# Major oscillations in spontaneous home-cage activity with an infraradian periodicity in C57Bl/6 mice housed under constant conditions

**DOI:** 10.1101/2020.09.09.290148

**Authors:** K. Pernold, E. Rullman, B. Ulfhake

**Affiliations:** Div. clinical physiology, Department of Laboratory medicine, Karolinska Institutet, CA

## Abstract

Using 14-20 months of cumulative 24/7 home-cage activity recorded with a non-intrusive technique and a data driven analytical approach, we here provide evidence for the existence of a circannual oscillation (1-2 SD of the mean, on average 65% higher during peak of highs than lows; P=7E-50) in spontaneous activity of male and female C57BL/6 mice held under constant barrier conditions (dark-light cycle 12/12 h (DL), temperature 21±1°C, humidity 40-60%). The periodicity of the season-like oscillation is in the range of 2-4 months (on average 97 days across cohorts of cages) and off-sets also responses to environmental stimuli but does not significantly alter the preference for activity during the dark hours of this nocturnal mouse strain (P=0.11 difference between highs and lows).

The significance of this hitherto not recognized slow rhythmic alteration in spontaneous activity is further substantiated by its co-variation with the feeding behaviour of the mice. The absence of coordination within and between cohorts of cages or synchronization to the seasons of the year, suggests that the oscillation of in-cage activity and behavioural responses is generated by a free-running intrinsic oscillator devoid of synchronization with an out-of-cage environmental time-keeper. Since the variation over time has such a magnitude and correlate with the feeding behaviour it is likely that it will impact a range of long term experiments conducted on laboratory mice if left unrecognized.

## Introduction

Organism behaviour shows distinct rhythmicities governed by internal oscillators which synchronizes (entrain to) external time-keepers (zeitgebers) such as light^1–4^. Most well characterized is the circadian rhythm (reviewed in ^5, 6^) but gravitational (tidal earth push and pull), seasonal, biannual and annual rhythms have also been described^7–12^. In brief, the free-running circadian rhythm (21-27h in small rodents) is generated endogenously by feedback loops involving expression of key clock genes expressed by neurons of the Suprachiasmatic nucleus (SCN) in the hypothalamus that synchronizes with the earth’s solar day (24 h). The entrainment of the circadian rhythm to a day occurs in mammals through retinal neurons directly responding to photic stimuli, or indirectly through retinal cones and rods, and is transmitted to several brain regions, importantly the SCN and, further, to the epithalamus. Here cells of the pineal body translate and feedback the message by supressing the conversion of serotonin (an amine that also serve as a major neurotransmitter in the nervous system) to melatonin^13, 14^ which is secreted during the dark hours only (*idem*)^9, 15–18^. The daily oscillation of the key gene products is not exclusive to the SCN where direct interaction occurs with the neuroendocrine (HPA) system within hypothalamus and key regions of the brain^17^, but is expressed in different peripheral tissues of our body^6^ explaining the circadian oscillators’ profound impact on bodily functions. A number of biorhythms have a higher frequency than the circadian and are collectively referred to as supraradian biorhythms. While some high-frequency rhythms are well established like the sleep-wake cycle and the different reoccurring sleep states, most supraradian rhythmicities have an unknown origin. At the other end of the spectrum we have the slow biorhythms (> 1 day), referred to as infraradian (tidal, lunar, seasonal, annual and bi-annual). Typical examples are the reproductive cycle, hibernation and seasonal migration to name a few. Irwing^1^ categorized slow biorhythms into being endogenous rhythms that synchronizes to an external time-keeper (zeitgeber) (type I), rhythms that are purely driven by an endogenous generator (type II), and those that are exclusively driven by re-occurring environmental changes interacting with the geno-/phenotype of a species such as seasonal allergic complications or the stress and behavioural deviations of small rodent responding to regular husbandry routines like the cage-change at an animal facility (type III).

Laboratory rodents like mice used in experiments are commonly bred and maintained under well-controlled and standardized barrier conditions. While husbandry routines provide light (dark and light period of the day; DL) to which the circadian rhythm entrains, these mice are not purposely subject to seasonal or annual variations in the environment. Albeit a few studies have presented data suggesting that both exploring and pain-related behaviours in small laboratory rodents show seasonal variations ^19, 20^, there is a prevailing scepticism towards the existence of seasonal/circannual behaviours in laboratory mice housed in constant conditions ^21^. This notion is building on the successful migration of the house mouse to almost every corner of the world attributed to the mouse capacity to adapt and reproduce in different environments across seasons^22, 23^. In line with this both the standardized care and use of laboratory mice commonly consider strain, age (seasons of the life-span), sex and the circadian rhythm as significant parameters but not seasonal (circannual) variations^22, 24^.

Behavioural assessments is still mainly done by bringing the animal out of its living quarter and social group context, and includes commonly also handling of the animal by the observer. As widely discussed, these experimental conditions may introduce serious biases invalidating the test result. The aspiration to obtain cumulative records of laboratory animal behaviours in a “non-intrusive home-cage like style” dates back more than a century (for an historical re-collection see^25^) but was until recently restricted to a limited set of tools like the running wheel integrated in the home cage (served critically for the understanding the circadian rhythm in small rodents). Over the past few years a number of techniques^26–40^ have become available for automatic non-intrusive 24/7 monitoring of in-cage activity^41^ of single housed^27, 32, 38, 42^ and group housed^26, 28, 29, 37, 39, 43–45^ small rodents. We recently explored data of cage floor activity of group-held mice using a DVC^™^ system to analyse daily rhythms of spontaneous in-cage activity^43^. In that study we noted a significant variation of in-cage activity across weeks during the 3 months of cumulative recordings at the different participating facilities; but within this time window we could not resolve any pattern of the oscillation in activity across weeks. Using the same DVC^™^ platform (Supportive informations Fig. S1) we here report recordings from 14 cages with male (n=11) and female (n=3) C57BL/6 mice starting at age 7-12 weeks until ∼16 months of age (n=11) and in three of these cages up to age ∼700 days (see Methods) with the purpose to reveal alteration of spontaneous in-cage activity over time.

## Materials and methods

### Mouse strain, sex and age

Cohorts of specific pathogen free (SPF, according to FELASAs exclusion list) male and female C57BL/6 mice were purchased from Charles River (strain C57BL/6J), Germany, or Janvier labs (strain C57Bl/6Rj), France, (Tables 1-2). Transported by car and upon arrival subjected to a brief health check, at 6-8 weeks of age, mice were randomly allotted to cages and grouped 4 or 5 per cage.

**Table I.**
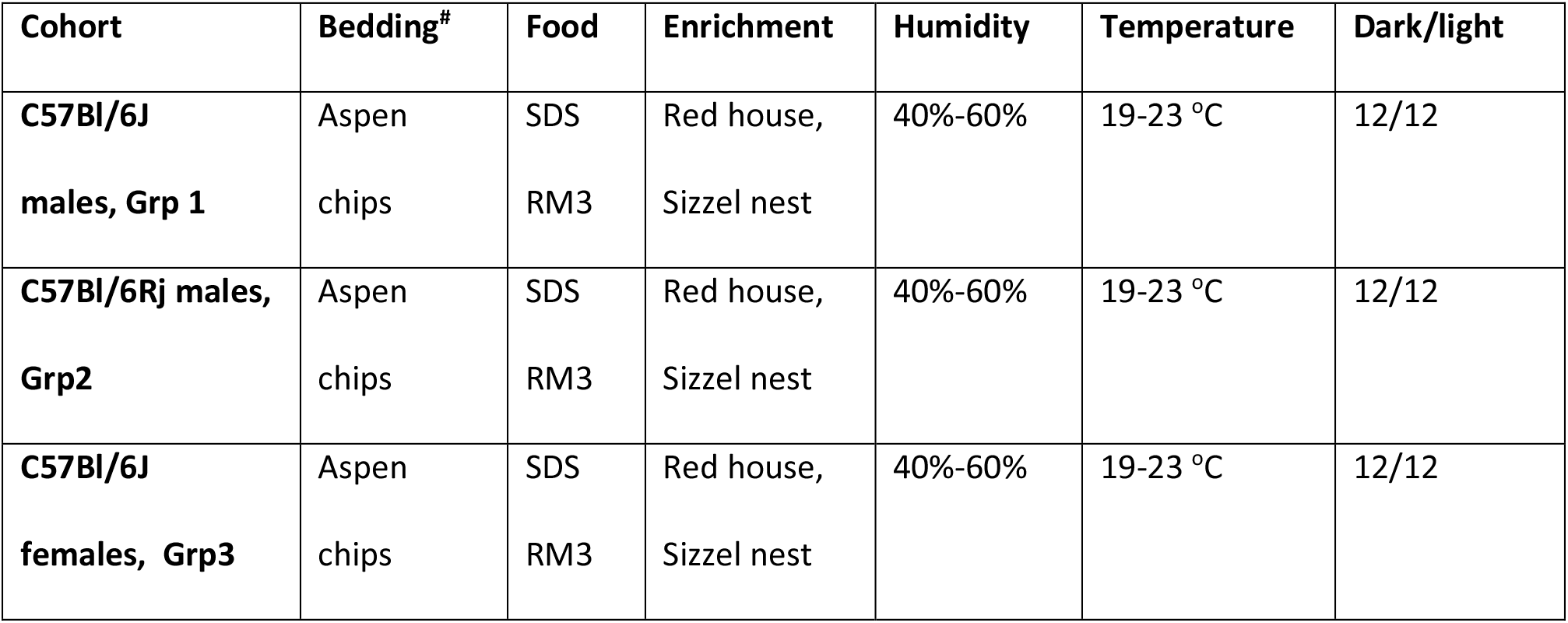

| **Cohort** | **Bedding<sup>#</sup>** | **Food** | **Enrichment** | **Humidity** | **Temperature** | **Dark/light** |
| --- | --- | --- | --- | --- | --- | --- |
| **C57Bl/6J**<br>**males, Grp 1** | Aspen<br>chips | SDS<br>RM3 | Red house,<br>Sizzel nest | 40%-60% | 19-23 °C | 12/12 |
| **C57Bl/6Rj males,**<br>**Grp2** | Aspen<br>chips | SDS<br>RM3 | Red house,<br>Sizzel nest | 40%-60% | 19-23 °C | 12/12 |
| **C57Bl/6J**<br>**females, Grp3** | Aspen<br>chips | SDS<br>RM3 | Red house,<br>Sizzel nest | 40%-60% | 19-23 °C | 12/12 |

**Table III.**

| **CC III**<br>**male (n=40)** | **Grp3**<br>**female (n=12)** | **Grp2**<br>**male (n=12)** |
| --- | --- | --- |
| **C57Bl/6J** | **C57Bl/6J** | **C57Bl/6Rj** |
| Charles River, Germany | Charles River, Germany | Janvier, France |
| Arrival date: 2016-Nov-22 | Arrival date: 2016-Nov-29 | Arrival date: 2016-May-10 |
| DVC – acclimatization 1(-2)<br>week – 5/cage | DVC - acclimatization 1(-2)<br>weeks – 4/cage | DVC - acclimatization 1-2<br>weeks – 4/cage |
| Weekly cage-change, no BW | Weekly cage-change incl. BW<br>and feed weighing | Weekly cage-change incl. BW<br>and feed weighing |
| DVC recordings started at<br>age 49 days | DVC recordings started at<br>age 50 days | DVC recordings started at<br>age 70 days |
| DVC recordings until age 483<br>days.<br>Record length is 434 days. | DVC recordings until age 476<br>days.<br>Record length is 426 days. | DVC recordings until age 699<br>days. Animals were out of<br>DVC during 64 days.<br>Record length is 565 days. |

### Holding and husbandry conditions

Mice were kept in GM500 IVC (Tecniplast SpA, Italy; type IIL) cages in a DVC^™^ system (Tecniplast) (see Supportive informations Fig. S1). IVC cages are individually ventilated with 75 HEPA14 filtered air exchanges per hour. Air temperature was 21 ±2°C, and air humidity 40-60%. The holding room had a 12-12h dark/light (DL) cycle with instant switch between light (white light 15-40 Lux inside the cage) and dark. Ad libitum access to food (SDS RM3 irradiated pellets) and weakly chlorinated water through water bottle that is changed every week. Cages had 100g aspen chips (Tapavei, Finland) as bedding, sizzle nest and red polycarbonate mouse house (Tecniplast Spa) as enrichment. The husbandry routines included weekly cage change (whole cage was changed but red house and some of the soiled bedding were moved along with the animals to the new cage) including body weighing and in two of the cohorts also weighing of feed on a fixed weekday (Tables 1 and 2). The holding unit was subject to health inventories according to FELASAs recommendation for a sentinel reporting system (i.e. the subjects of the study was not directly affected in the health inventory) four times a year^46, 47^ and during the study period the output from the sentinel system in use met the FELASA exclusion list for specific pathogen free animals (SPF). Surveillance of health included daily check-ups and weekly individual examination during the cage-change and weighing. As needed the designated veterinarian of the facility was consulted.

### Ethical considerations

The experimental procedures were agreed upon, reviewed and approved by the Regional Ethics Council, Stockholms Regionala Djurförsöksetiska nämnd; project licenses N116-15, N115/15 and N184/15 plus amendments.

### DVC recordings

In total recordings were collected from 14 cages maintained at the facility for an extended period of time and belonging to 3 different groups (Table 2). Group 1 (CC III) with 8 cages each populated with 5 male mice and this group is age- and season matched with group 3 having 3 cages with 4 females in each cage (Table 2). Group 2 with 3 cages housing 4 male mice per cage arrived at the facility about 6 months earlier than the two other groups (Table 2).

The core of the DVC system is an electronic sensor board installed externally and below each standard IVC cage of a rack (Fig. S1). The sensor board is composed of an array of 12 capacitive-based planar sensing electrodes. A proximity sensor measures the electrical capacitance of each of the 12 electrodes 4 times per second (i.e. every 250 msec). The electrical capacitance is influenced by the dielectric properties of matter in close proximity to the electrode, leading to measurable capacitance changes due to the presence/ movement of animals in the cage above. Thus, movements across the electrode array are detected and recorded as alterations in capacitance (Fig. S1). By applying custom designed algorithms to the collected data we can infer information regarding in-cage animal activity^43^. For this study, we used the first-order difference of the raw signal (i.e., capacitance measured every 250ms) as the basic metric of animal activity. More specifically, we take the absolute value of the difference between two successive measurements for each electrode (signals spaced 250 msec apart) and compare it against a set threshold (capacitance variations due to noise) to define an activity event. This metric thus considers any animal movement that generates a significant alteration in capacitance, an activity event^43^. In this study we have used the average number of activations across all 12 electrodes per minute, i.e. the average number of activation events across the 240 time-slots per minutes of all twelve electrodes (for examples of activation plots see Supportive information Fig. S2). For further details on use of the spatial resolution provided by the electrode array, please see ^43 48^. Note, this activity metric represents the overall in-cage activity generated by all mice in a cage from any electrode and is not tracking activity of individual group-housed animals (for activation plots of the cages used in this study see Supportive information Fig. S2A-N). However, as shown elsewhere there is a close correlation between the metric activations used here and locomotor activity of the animals in the cage^48^. Moreover, the DVC^™^ records activity on the cage floor only.

### Data processing and statistics

Data were processed through scripts in R (version 3.5.0). We used spectral analysis, functional data analysis, linear regression and statistical analysis of longitudinal data; all scripts are open source and available for R.

The data file from each cage (see below for data availability) was plotted with minute resolution as activation plots (Supportive information Figs 1-2) or heat maps (see Fig. 1 A, C) to visualize variations in activations across day and night (circadian rhythm; Fig. 1A), and over time (Figs. 1C). Heat maps were constructed using ‘ComplexHeatmap’ script (version 2.0.0) available for R^49^. By linear-regression across the whole recording period, the over-all change in activity was assessed for each cage and expressed as coefficient of determination (r^2^) with level of significance (Figs. 2 and 4, and Supportive information Table S1)).

**Fig. 1.**
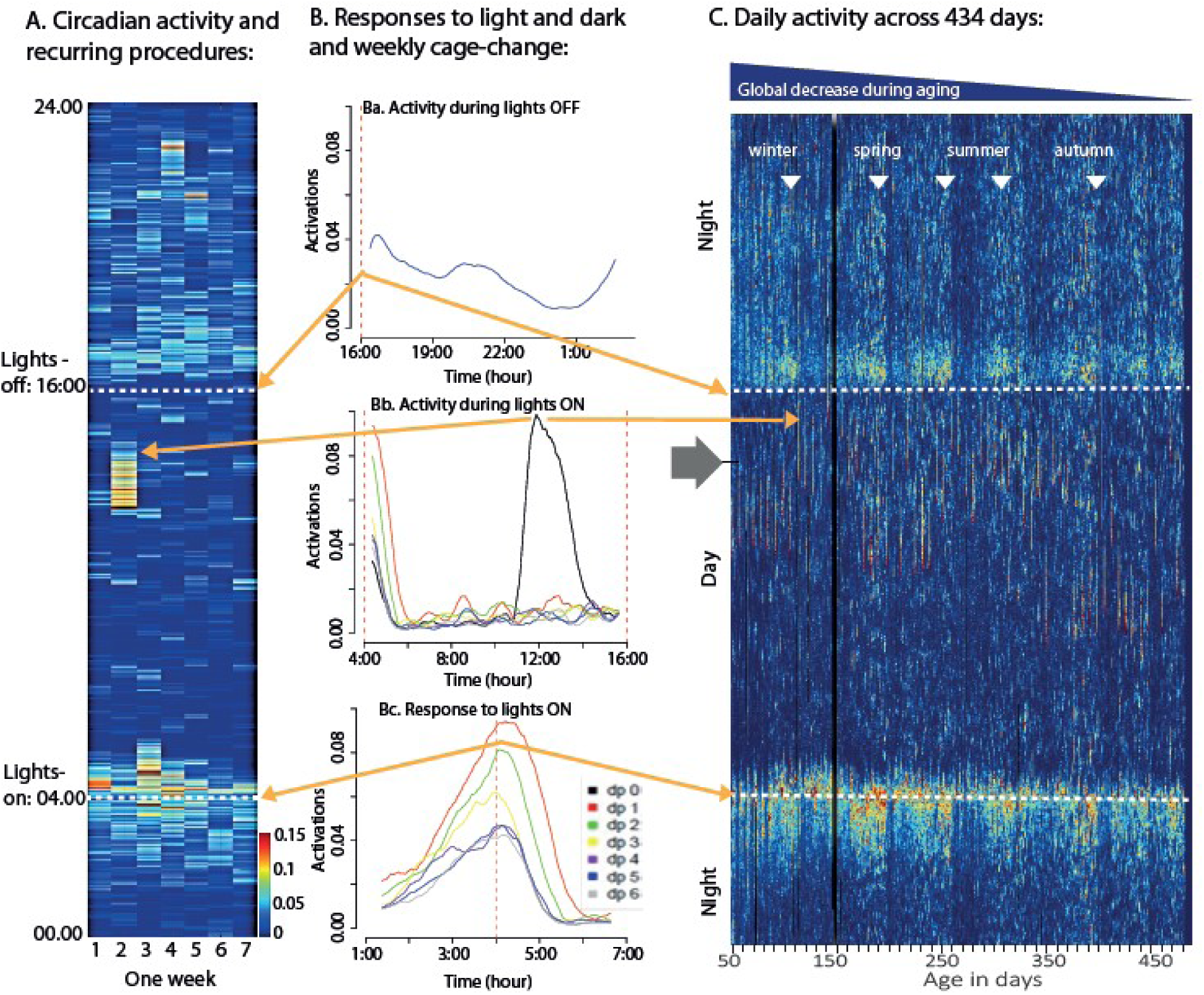
(A) To the left is a heat map with seven columns corresponding to the days of one calendar week (day 1 to 7) with colour coded activations (for conversion see scale to the right) illustrating the daily rhythmicity in activity synchronized to lights-on (4 AM, white stippled line in A and C) and lights-off (4 PM, white stippled line in A and C) in one cage (A04) with 5 male mice. The details of the response to lights-on is shown for each day in Bc lower graph left of the heat map (separate colour coded traces for each week day, with dp0 being the day of the cage change (see key to the right), averaged across weeks); followed in Bb by the day-time resting pattern which is regularly interrupted by minor bursts of activity (middle graph; each day is colour coded as in Bc). According to the standard husbandry routines the cage is changed once a week (day 2 in A and dp0 in graph) which instigates a dramatic increase in activity lasting for several hrs and with a carry-over impact on the response to lights-on the next following days (see Bc). In response to lights-off this nocturnal species increase activity reaching a peak within 1h (Ba). During late night-time activity decreases towards the resting pattern of day-time until the response to lights-on is triggered. In (C) the whole data-set of activations across the 434 days (see Table II) is shown for the same cage (A04). The ordinate is the same as in (A) while the abscissa is animal age in days (49 to 483 days of age). Deep blue indicates no activity or data missing. The elements of the circadian rhythm across day and night in (A) is also evident in (C) (orange arrows to aid navigation). Across this phase of the mice life-span there is an over-all decrease in activity (blue triangle on top of panel C) which is small but yet statistically highly significant (−4% during 400 days for this cage; P=2E-10; see also running text and Supportive information Table S1). Inspections of the long term records of activations (C) revealed a marked slow oscillation of peaks (highs; indicated by filled inverted triangles) with intervening lows.

**Fig. 2.**
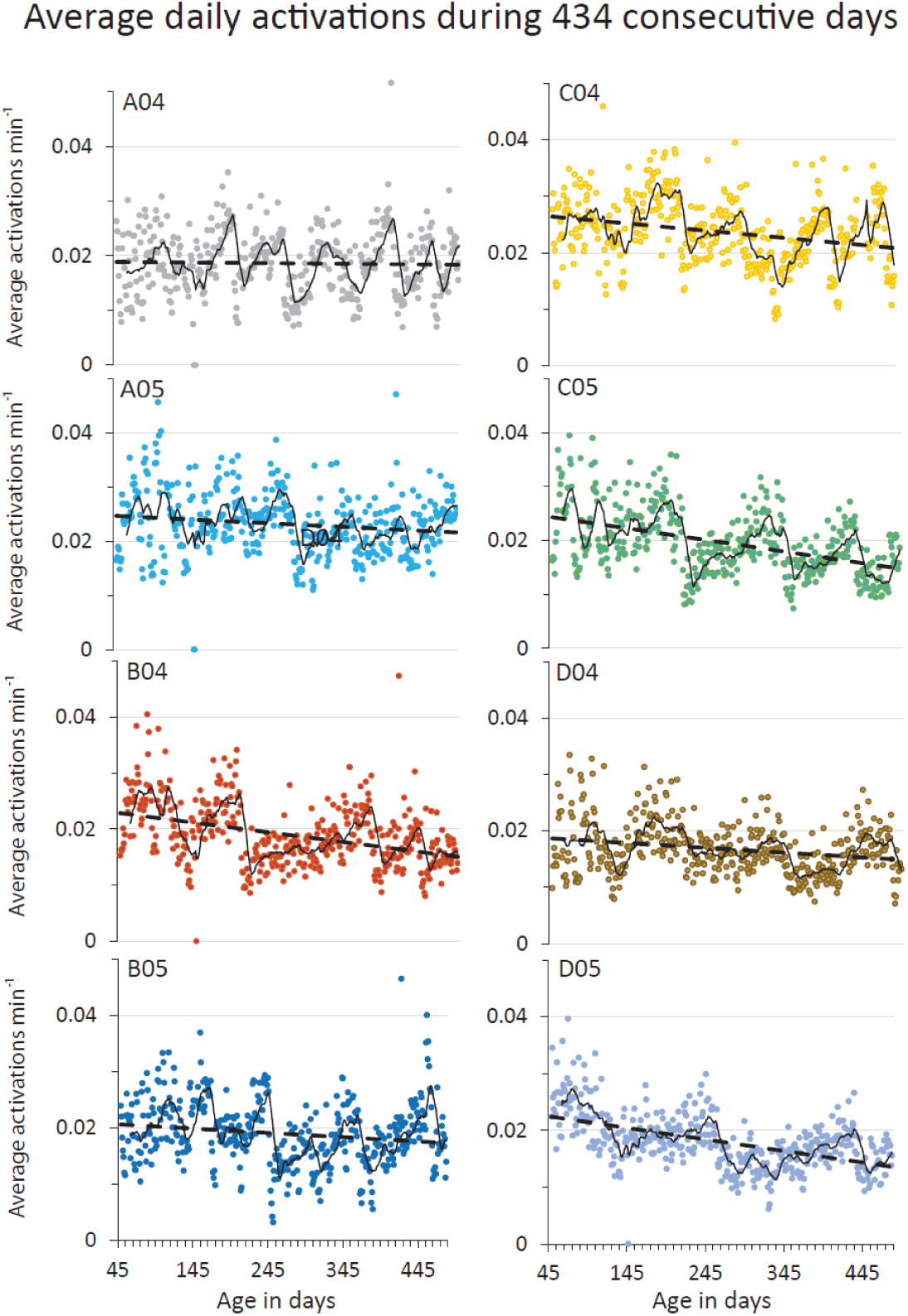
Plot of average daily activations for each of the cumulative data sets in Group 1; A04 depicted also in Fig. 1 is the top left panel. Cage ID is in each panel in the top left corner. Evident in all plots is that the overall activity decreases with age as indicated by the interrupted black line in each panel (for linear regression details see Supportive information Table S1). The continuous black line of each panel represents the moving average using a window of one week. As evident from the distribution of data points in each panel and the moving average trace, highs and lows are present in all cages but the dates for highs and lows are not synchronized across cages (for cage A04 the inverted triangles in Fig 1B are the first five peaks). The data is rather noise during the first weeks which we attribute to the acclimatization of the animals to new groups and cages. Nevertheless, in each panel three to five peaks with intervening lows can readily be identified. Ordinate is average activation min^−1^ during one day and night, i.e. 24h. Abscissa at the bottom of each column is valid for all its panels and shows the age of the animals.

**Fig. 3.**
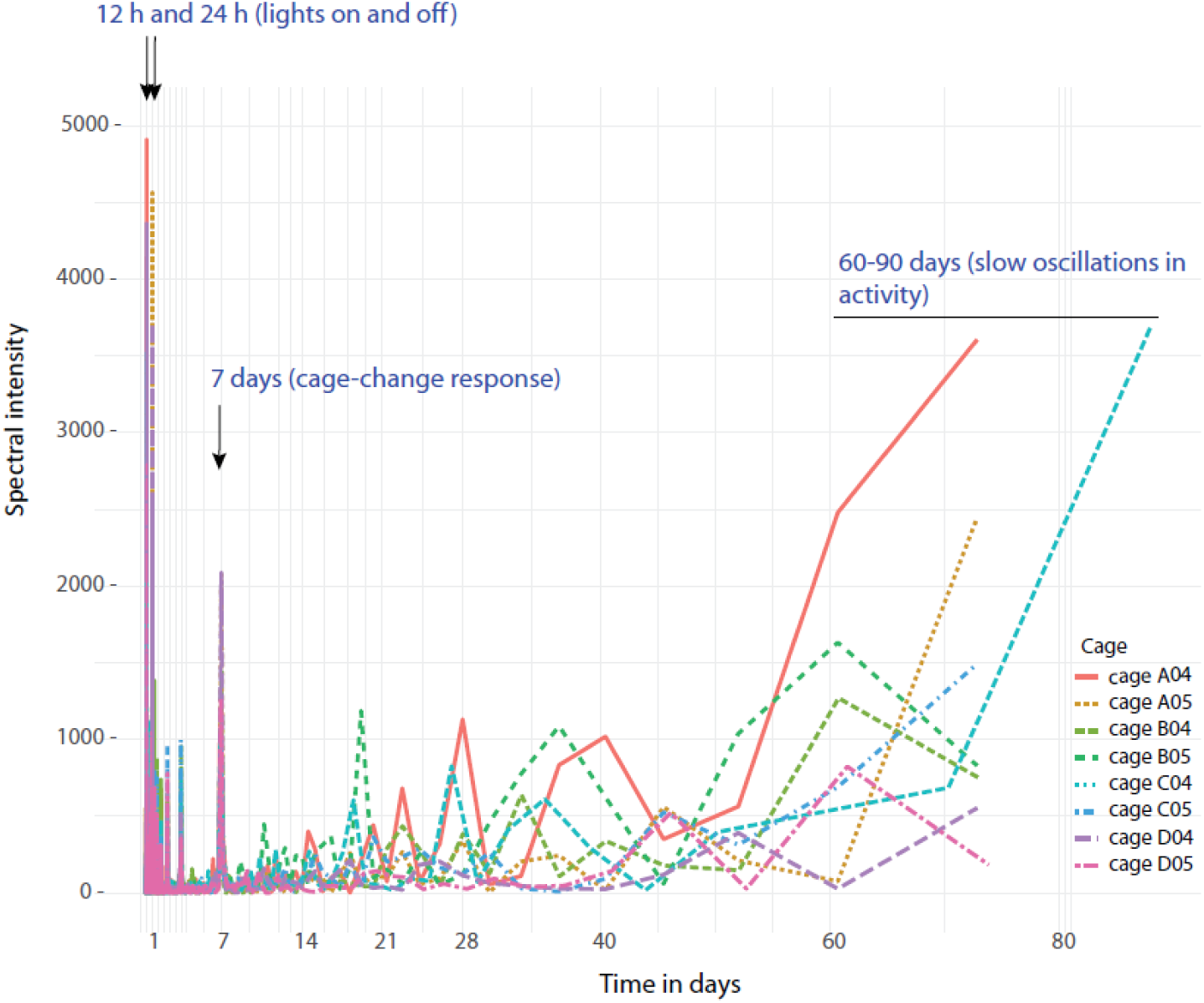
Superimposed spectrograms of activations clustering at different frequencies of the eight cages (colour coded; key is to the right in the spectrogram) with male mice in group 1 (see Methods). Note the precise timing across cages of the behaviors triggered by environmental stimuli, while the slow oscillations having a similar periodicity are not synchronized across cages or with seasons (Fig. 4). The slow peaks are generated by only 3-6 instances in the cumulative data records while e.g. lights-off occurs 434 times in each of these data sets.

**Fig. 4.**
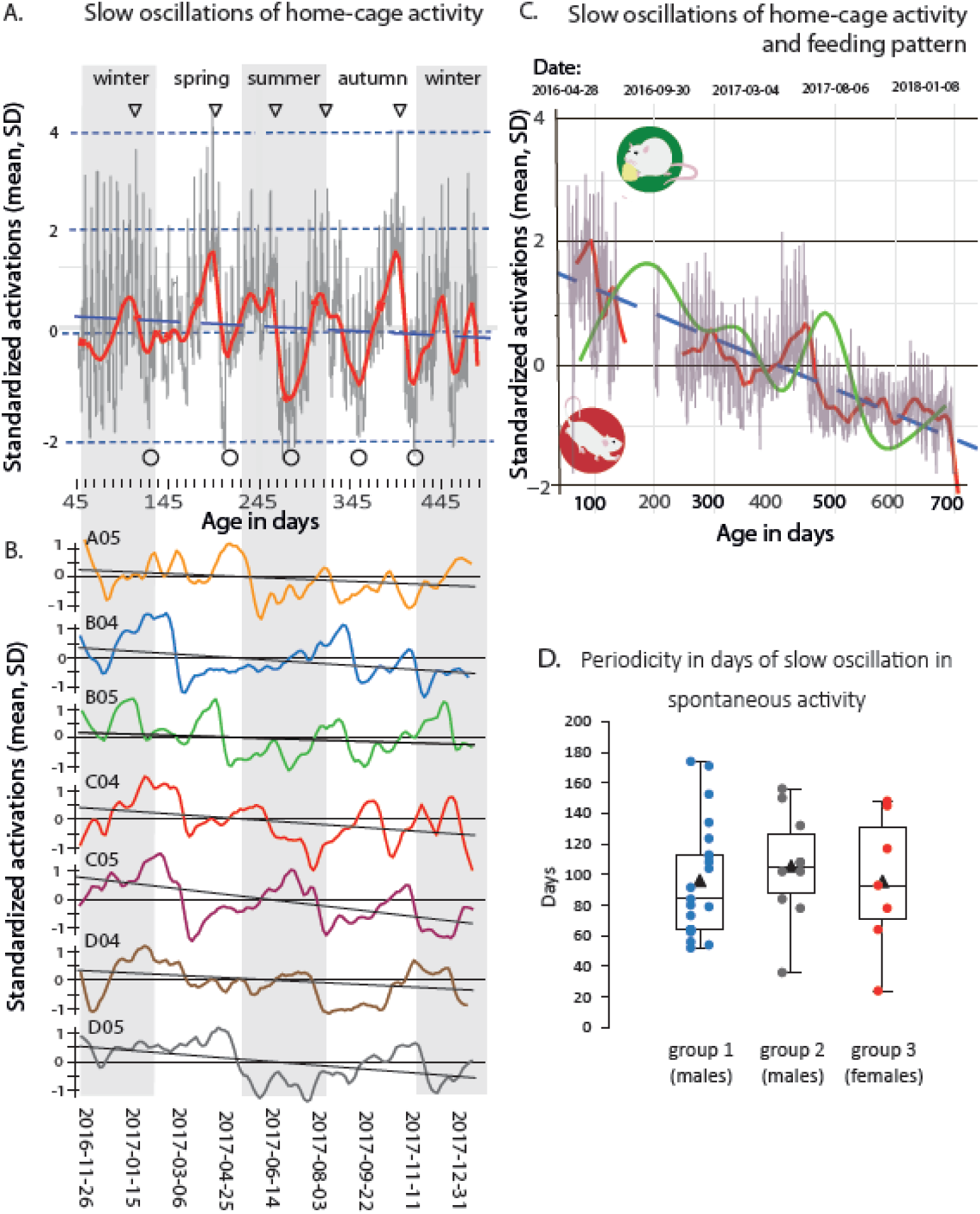
**A** Upper panel shows plot of normalized (see Methods) activations (ordinate is mean and standard deviations of the mean) for the data set shown in Figs. 1C and 2, with highs indicated by inverted triangles (same as in Fig. 1C) and lows by open green circles at the bottom. On top of the slow rhythm and having a similar amplitude is the recurring response to the cage-change which occurs at regular interval of one week (for further details of the plot see Supportive information Fig. S3). The slow rhythmicity is indicated by the red curve and the interrupted blue line is the linear regression showing the over-all decrease in activations with advancing age. The data has been plotted against dates (abscissa bottom), age in days (abscissa middle) and with the seasons of the year as background (abscissa at the top, intervened grey-and-white vertical stripes). **B** Lower panel is the slow rhythmicity (continuous line with different color for each cage) of the complementary 7 cages in Group 1 along with the linear regression of the each cumulative record showing the decrease with advancing age (black line). The annotations on the abscissa (season, age in days and dates) are the same as in A. **C** shows normalized activations (as in A and B) up to an age of 699 days in one cage belonging to group 2 (A07; activations data available in Supportive information Fig. S2I) where food consumptions (average across one week; see Methods) was measured (green line). The red curve indicates the slow oscillation in activations (stippled line indicates when the cage was not in DVC; see Table II) while the interrupted bluet line is the regression of overall change in activity with advancing age. The slow oscillations in food consumption (green) and activations (red) are highly correlated r^2^=0.72. **D** Boxplots visualising the period i.e. peak-to-peak of slow oscillations in undisrupted home-cage activity of the three groups of cages (group 1, male mice, housed 5 to a cage; grp2, male mice housed 4 to a cage; and grp3, female mice, housed 4 to cage). Box indicates median, 25%-75% quartiles with max and min as bars. Filled triangles indicate mean periodicity of each group. The observation from each cage in each of the groups have been indicated by separate colour-filled circles. The average period is similar in group 1-3 (96, 106 and 96 days, respectively).

Spectral analysis and periodograms were generated using the periodogram-function from the ‘Time Series Analysis’ (TSA) library v 1.2^50^. Minute-averaged activity data (across electrodes and samples min^−1^) was used and each cage was analysed separately. In order to account for the small but consistent decline in activity over time and make the time-series stationary, a linear model was fitted (Activity∼ Time) for each time-series where the residuals of the model was used as de-trended times-series. Stationarity was confirmed using Augmented Dickey-Fuller Test where a p-value of <0.01 was considered a stationary time-series. The resulting periodogram of the de-trended activity-data was plotted with wavelength (i.e. time between peaks) on the x-axis and spectral intensity on the y-axis (Fig. 3). In the same fashion, spectral intensity as a function of time was analysed for each cage using the Spectro-function from the ‘seewave’ library v 2.1.4^51^ with a window-length of 7 days (10 080 minutes) and a smoothing overlap of 25%. To improve readability, the output was divided into three separate graphs covering wavelengths 0-12hours, 12-240 hours and 240 hours (10 days) to Infinity. Pre-set graphical parameters were used where red-green-blue-black denote higher to lower spectral intensity.

For the generation of smoothed time-series, minute-averaged activity data from each cage was centred and scaled to a mean of 0 with standard-deviation as principal unit where 1 is activity 1 standard-deviation above the average activity throughout the observation period. Smoothing was achieved by applying a 24 hour rolling mean followed by a 5% loess-regression using the loess Fit-function from the ‘Linear Models for Microarray Data’ (limma) library v. 3.42.0^52^. Graphs were thereafter generated by plotting 24 h averaged activity over time together with the smoothed activity data (Fig. 3A and 3C and Supportive information Fig. S3).

The peak response to lights-on^42^ was expressed as the highest number of activations min^−1^ minus base-line activity (see below) during 180 minutes of recordings prior and subsequent to the time at which lights-on took place^43^ the day after cage-change (Figs. 1Aa and 5). Records with missing data (exceeding 59 min) or any procedures, or any other non-regular holding unit event interfering with the recording period of 360 minutes, were excluded.

Like-wise the peak response (activations min^−1^ minus base-line activity min^−1^) to lights off covered records 120 minutes prior to and subsequent to time of lights off and is the average peak response during a cage-change cycle at a high or a low (Figs. 1Ac and 4). Exclusion criteria were the same as for lights on.

Day and night time activity, respectively, was expressed as the average activations min^−1^ during the period with lights-on and lights off, respectively, while daily activity is the average activations min^−1^ across 24 h. Weekly average activations per minute is the corresponding value calculated for seven days (= cage-change cycle) and used to express base-line in-cage activity during weeks of highs and lows, respectively (Fig. 5 and below). The fraction of the daily activity occurring during lights-off was calculated from these metrics (Fig. 6).

**Fig. 5.**
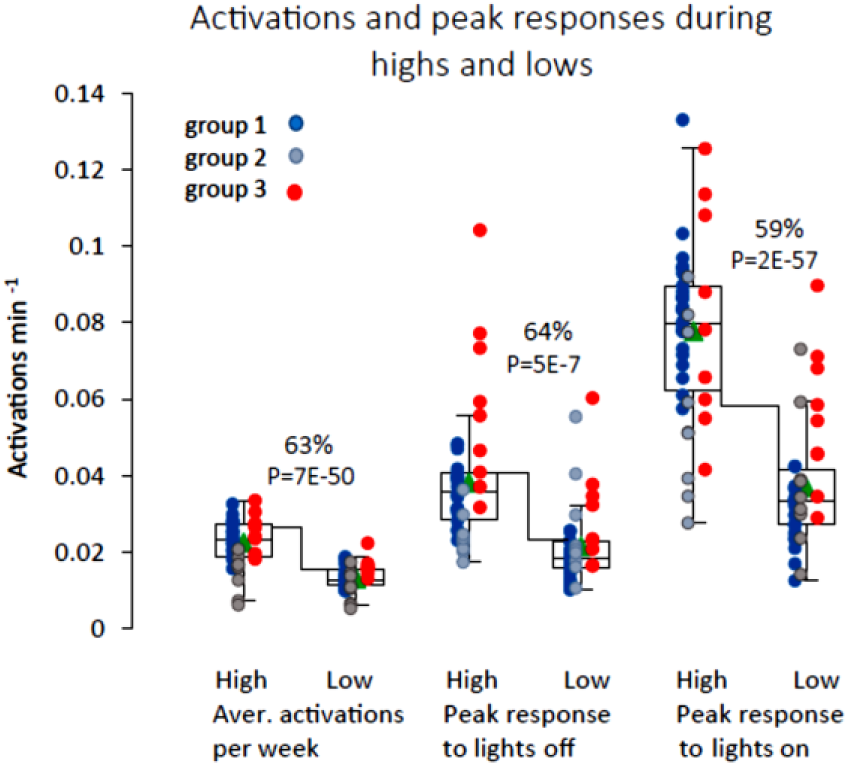
Boxplots of weekly average activations during weeks of highs and lows, respectively (63% difference between highs and lows, P=7E-50). Cages belonging to the different groups (Methods) have been colour coded; blue is group 1 (males, five to a cage), grey is group 2 (males, four to a cage) and red is group 3 (females, four to cage). Peak response in activations (activations above weekly average) instigated by the events lights-off (left) and lights-on (right) for the three groups of cages during weeks of highs and lows. The differences in the responses to lights-on (+59%) and lights-off (+64%) between weeks of highs and lows, respectively, are significant (P=5E-7 and P=2E-57, respectively) and have a similar magnitude. The amplitude of both responses are closely scaled in size of the change of the underlying difference in average weekly activity.

**Fig. 6.**
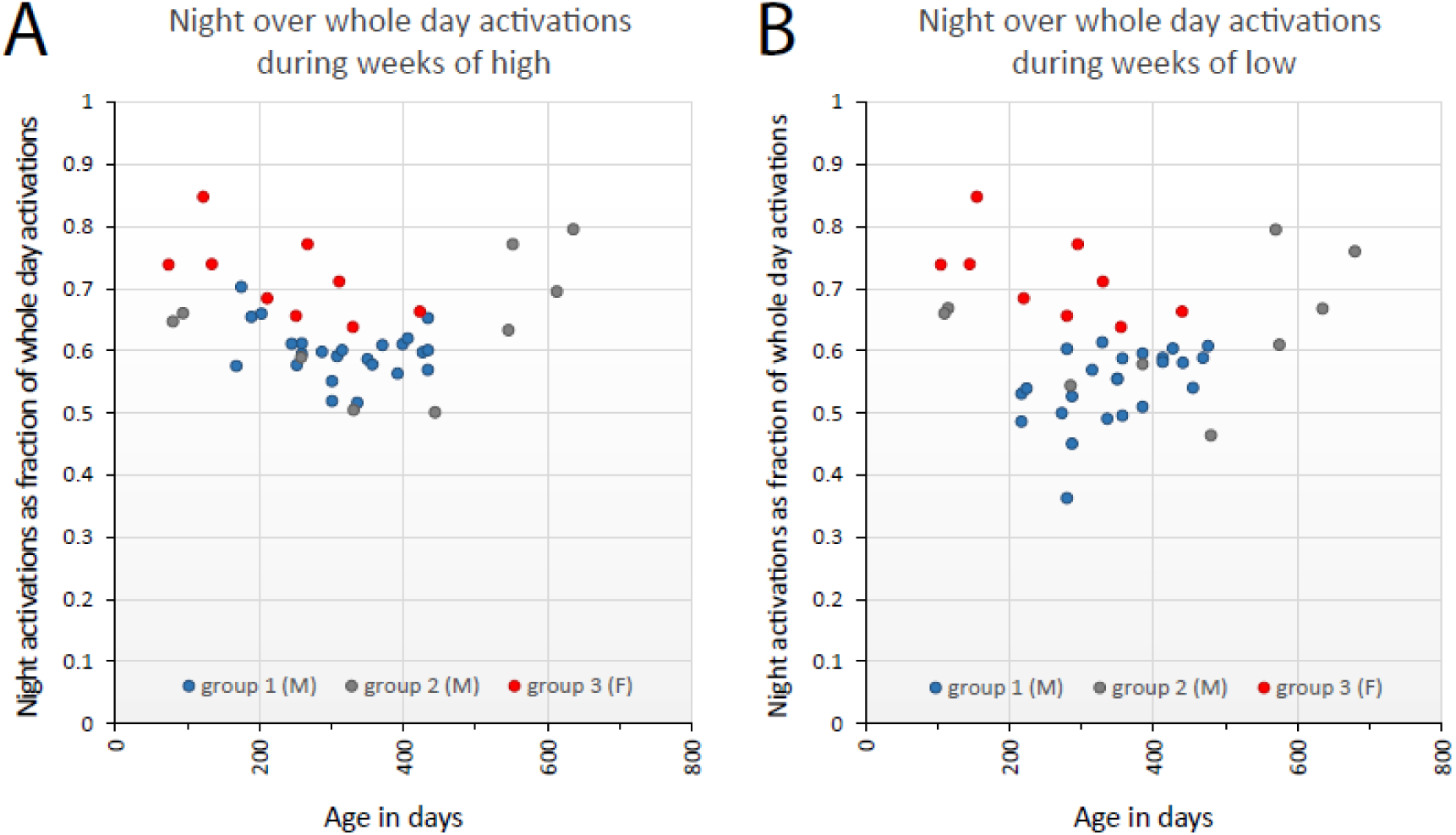
Plots of the fraction of the daily home-cage activity that occurs during the dark hours (night) over weeks with high (left panel; average 63%) or low (right panel; average 60%) level of home-cage activity, respectively. Ordinate is fraction of total daily activations occurring during dark period and abscissa indicates age in days of the animals. The cumulative data records covers 434 days for groups 1, 426 days for group 3, and 565 days for group 2. Key to cohort group at the bottom of the graphs. There are clear difference between cages within and between groups of cages but also a considerable variability over time of each cage. Typically two-thirds of the total daily activity occurs during night time both at highs and at lows and there is no significant difference between highs and lows (P=0.11) or trend of change over time (P=0.10). Thus, the circadian rhythm of these nocturnal mice are essentially preserved between highs and lows. In contrast there is an impact by sex (P=2E-10), females allotting more of the daily activity to the dark hours.

We used the rank-based analysis of variance-type statistic (ATS) to test differences within and between cages and sex ^53^. Analyses were implemented using the nparLD package in R statistical software ^54^. npar-LD is a non-parametric test for longitudinal data that does not require strong assumptions as the Repeated Measures ANOVA ^43^. We considered cages as subjects, high-or-low and observation-order as within-subject factors (“sub-plot” repeated factors), and sex as between-subject factors (“whole-plot” factor). According to authors’ terminology ^54^, we used F1-LD-F2 (sex as the whole-plot factor and with high-or-low and observation order (age) as the repeated ones). Since high-or-lows are arranged chronologically the observation-order and interpreted as the age-factor.

## Results

The DVC output record (average number of activations across all 12 electrodes min^−1^) of each cage was first plotted as a heat map with full resolution (Fig. 1A and C). Inspection of these heat maps revealed the circadian rhythmicity in activity synchronized by lights on (Fig. 1Aa) and lights off (Fig. 1Ac). In addition, the response to the weekly cage-change was readily identified (Fig. 1Ab)^43^. Closer inspection revealed a significant decline in over-all activity over time (Figs 1C and 2; see also Fig. 4A-C). On average, the daily activity changed by −18% (group 3), −22% (group 1) and −28% (group2), respectively, from age 70 to age 470 days. As evident from the slopes of the linear regression (Supportive informations Table S1) of the cumulative records of activity for each cage, there was a large cage-to-cage variability in the age-associated decline in activity (from <-1% to −43%) without any corresponding clinical sign of compromised health or welfare among the animals.

An even more conspicuous feature of the heat maps (Figs 1C) and activation plots (see Supportive information Fig. S2 A-N) is the recurring periods with higher over-all in-cage activity (‘highs’; triangles in Fig. 1C intervened by periods with lower over-all activity (‘lows’). This pattern of oscillation of in-cage activity was present in all cages examined and Fig. 2 depicts the eight cages of group 1 showing the magnitude of the activity-shift between highs and subsequent lows (for plots of data with minute resolution of all 14 cages please see Supportive information Figs S2 A-N). On average, the difference in activity between the peak of a high and the subsequent low was 1.6 (±0.61 SD) standard deviations of the mean.). Both the heat maps and activation plots of in-cage activity suggested that activity cluster at different frequencies possible to resolve by spectral analysis and functional analysis.

Using spectral analysis, the full data-set with minute resolution of each cage was analysed considering the level of activations with different peak-to-peak periodicity and plotted as a periodogram (Fig. 3). From these plots it is evident that high levels of activity peak around certain expected frequencies such as lights-on and lights-off (circadian rhythm) at 12 and 24 hrs (Figs 1 A, Ba and Bc) and at 168 hrs (7 days; Fig. 1 A and Bb) which is the instant impact on activity by the weekly cage-change. In addition we observed both low-amplitude high-frequency peaks of unknown origin and a high amplitude cluster with long wave-length (Fig. 3). The peaks associated with the identified external triggers and several of the low intensity high-frequency peaks are synchronized across cages (Fig. 3) while the high intensity low-frequency cluster seems to be asynchronous across cages (Figs 3, see also Fig. 2 and Fig. 4 A-B).

To deepen the analyses of the low frequency oscillations in spontaneous in-cage activity, we generated smoothed time-series centred on the mean of the minute-averaged activation data for each day and cage (see Material and methods) and plotted daily averaged activity together with the smoothed activity data against, age of the animals, calendar date and season of the year (Fig. 4 A-B; see also Supportive information Fig. S3). The calendar dates of the slow-frequency highs and lows can be extracted directly from the plots (or by the script) and shows a periodicity in the range of 2-4 months, with an average of 97 days for all groups of cages in this study (Fig. 4D) and, furthermore, that the slow oscillations in activity level are not synchronized between cages or locked to seasons of he year albeit having a frequency of about 4 cycles per year (*idem*).

Using the calendar dates of highs and lows, we calculated that the weekly average in-cage activity during highs was 63% higher than during weeks of lows (p=7E-50, Fig. 5). We also confirm that female mice have a higher in-cage activity than males (P=1E-3 and interaction sex*highs-lows P=2E-3; see Methods). Comparisons of the responses to lights-off and lights-on during highs and lows (Fig. 5) revealed that response amplitude was scaled to the difference in the over-all activity and during weeks of highs it was 64% and 59%, respectively, above the response during lows (P=5E-7 and P=2E-57, respectively). Thus these responses are more vigorous in terms of peak activations during highs than lows (*idem*).

Laboratory mouse strains are considered nocturnal^55–57^, i.e. show the greatest activity during the dark hours. However some mouse strains have a preference for being active day time, i.e. they are diurnal^58^, and there is evidence that nocturnality vs diurnality may at least to some extent be context dependent^4^. Commonly used laboratory strains like the C57Bl/6 are under facility housing conditions nocturnally active while day time is the main period for resting^43^. Here we analysed the proportion of total daily activity that occurred during the dark hours on a weekly basis and could not detect any consistent difference between weeks of highs and lows (P=0.11), respectively, or over time (P=0.10); 63% (highs) and 60% (lows) of the total daily activity occurred during the dark hours (Fig. 6). In contrast, there was a significant impact by sex (P=2E-10), with females showing a stronger preference for activity during the dark hours (72%) than males.

In 6 of the 14 cages analysed we had access to food consumption (see Materials and methods) and the ad libitum feeding behaviour also show a recurring variability over time having a periodicity similar to the slow oscillation in spontaneous activity. As depicted in Fig. 4C there is a close covariation between these two metrics with the increase in feeding following upon a period with increased activity.

## Discussion

Using DVC^™^ records of cage-floor activity extending over more than one year of group-held female and male C57BL/6 mice, we provide evidence for a slow season-like oscillation in over-all activity with an amplitude of about 1-2 standard deviations of the mean and a periodicity on average of 97 days, range: 2-4 months (Figs 1B, 2-4; and Supportive information Fig. S2A-N). We show that spectral and functional time-series analysis using open source applications (see Materials and methods) are useful approaches to characterize in-cage activity and we envisage that this approach can be elaborated further and expanded to time-space since the DVC^™^ provide a spatial resolution as well (see also^59^). Our results also confirm previous data that female mice of the C57Bl/6 strain are more active the male mice^43^ and add that a further sex difference is that females allocate a significantly larger fraction (72%) of the daily activity to the dark hours than male mice do. Still both sexes are nocturnal with about two-thirds of the activity during night time and the season-like oscillations in spontaneous activity described here did not alter the preference for being nocturnally active and to rest during day time (Fig. 6).

Spectral analysis of activity clustering with different periods (Fig. 3) revealed a number of supraradian frequencies with major peaks at lights on/off every 12 h followed by the circadian rhythm at 24 h (*idem*). High frequency rhythms was often synchronized across cages signalling that they had a common external time keeper/trigger. We do not know the origin of most of these short frequency clusters yet but by using spectral analysis and time series combined with more detailed annotation of parameters in the holding room (staff mobility, cages in/out of the rack, vibrations, noise, spectral or intensity fluctuation of the illumination, etc.)^16, 17^ we may be able to map their origin in the future. For example, short bursts of recurring activity during daytime in between periods of rest may be a trailing effect, where vibration and noise from one cage trigger activity in the next cage and so on; such patterns being close to synchronized may be possible to track down with the DVC recording technique. In the long wave-length spectrum, the synchronized response to the weekly cage change and the slow season-like oscillations was the peaks with the highest amplitudes (fig. 3). The latter is not synchronized across cages and as shown by the time-series analysis (Fig. 4) disclose a variability in period both within and between cages (see also below). Because of this and that these clusters occur only 3-5 times in the cumulative record of each cage, the spectral analysis could not determine the period of the slow oscillation with any precision except for having a period longer than 60 days. As indicated by our time series analysis the period is in the range of 2 to 4 months with an average period around 100 days. This slow –season like-oscillation in spontaneous activity was recorded from cages with male and female C57Bl/6 mice maintained under constant conditions (Material and methods) and is of unknown origin. Visual, sound/vibrations or olfactory cues could pass the barrier protection under which they were housed and signal seasons or other recurring alterations in the facility environment. However, the lack of synchronization between cages or to the seasons of the year (Figs. 2 and 4; see also supportive information’s Fig. S2). suggest that the slow oscillation in daily activity is intrinsic to the group of mice in the cage and may be a behavioural trait of this laboratory mouse strain^1^. As such it should be taken into consideration in a range of experiments were mice behaviours are used as bio-assays^60^ and may also be a metric of value in the context of animal welfare surveillance. Mice are group-living, social creatures thus it is not surprising that behavioural rhythmicities of daily activity is coherent among the group members especially since their living quarters are confined to the cage^4, 33, 61^. Even though we have no evidence, this seasonal-like rhythmicity could either be an endogenous oscillator which do not entrain to an external time keepers (type II rhythm ^1^) or a “free-running” endogenous generator lacking external trigger to synchronize with (type I rhythm ^1^). In favour of the latter is the variability in period (2-4 months) of the slow rhythmicity in spontaneous activity which bears resemblance to the variation of the circadian rhythm when being devoid of external stimuli to synchronize with (∼21-27 h c.f. 24h of the earths solar day).

Seasonal variation is a common phenomenon in natural settings of mammalian and non-mammalian species and includes among other things reproduction cycle, hibernation, migration and moulting ^1, 3, 9, 62^. There are evidence showing that seasons may be internalized by photoperiodic stimuli^3, 9, 63^. Thus, difference in photic stimulus (longer/shorter nights) over seasons impacts the production of melatonin in the pineal body so that with increasing length of the dark hours, the duration of melatonin secretion increases and vice versa^9, 15^. “To know in advance” is an important capacity for survival in the wild, for example foraging in advance of seasonal hibernation. The free-running circadian rhythm (21-27h in small rodents) is generated endogenously by feedback loops involving mainly 10 gene products (Clock, Bmal1, Perl 1-3, Cry1 and 2, Ror-α, −β and −γ,)^5, 6^ expressed in neurons of the Suprachiasmatic nucleus (SCN) that synchronizes with the earths solar day (24 h). The entrainment of the circadian rhythm to a day occurs by supressing the conversion of serotonin to melatonin^13, 14^ which is secreted during the dark hours only ^9, 15, 56^. While mice of the C57Bl/6 strain expresses all the clock genes in the SCN and peripheral organs it carries a loss-of-function mutation in the arylalkylamine N-acetyltransferase gene, encoding a rate-limiting enzyme in the conversion of serotonin to melatonin^64–66^. Thus the common but not exclusive mechanism to decode earth’s solar day or seasons through photoperiodic stimuli does not work in C57Bl/6 mice. Nevertheless, these mice respond to lights-on and lights-off and as shown elsewhere phase-shift of the circadian rhythm can also be induced by non-photic stimulus^67^.

The global impact of the circadian rhythm on peripheral tissues is mediated through secreted melatonin, endocrine signals and the autonomic nervous system^5, 17^. In contrast to the circadian rhythm, the onward signalling to the target organs (ovary and testis in the case of reproduction) of circannual rhythms are not fully understood. The TSH (thyroxin stimulating hormone; subunit B)-and Tx (Thyroxin; T3 and T4) axis via the thyroid hormone activating (Dio2) and deactivating (Dio3) enzymes, is one candidate system that has attracted interest^68–70^, and experimental evidence supports a link between melatonin secretion and circannual oscillations of this axis (idem). Further, TSH-Tx axis has the capacity to impact gonadal hormones as well as growth and metabolism of peripheral tissues. Although adaptation to seasons often appears to make use of the time-keeping machinery of the circadian clock i.e. photoperiodic stimuli converts to changes in the production and secretion of melatonin, the seasonal variation in the TSH-Tx axis is independent of the clock gene Per 2^70^. Moreover, for non-mammalian species infraradian rhythmicities independent of the circadian clock machinery have been discovered^10, 11^. Of interest to our observed correlation between feeding rhythm and slow oscillation in spontaneous activity, is the food-anticipatory-activity (FAA) generated by the food-entrained-oscillator (FEO). FAA rhythmicity has been shown to be independent of the Per 1-3 clock genes and suggested to use a different time-keeping mechanism than the circadian clock^71^. Taken together there are evidence suggesting that infraradian biorhythms can depend on, or be independent of, the time-keeping of the circadian clock.

Our data, including the covariation of the amplitude of stimuli responses with the general activity level during highs and lows (Fig. 5), provide unbiased support to earlier observations indicating that behaviours show season-like (circannual) variations in laboratory mice kept under constant conditions^19, 20^. In addition, there are also variances in mice behavioural responses within the phases of the circadian rhythm^72^ and infraradian variations in activity linked to the circadian circuitry^44^. Taken together, rhythmicities need to be considered in behavioural assessments used as the high-end readout of organismal function and in other bioassays of disease models and interventions, as well as in phenotyping screens^60, 72–76^.

Strains originating from the house mouse (*mus musculus*) are currently the most widely used mammalian animal models in experimental research, a main reason being that strains of this species were very useful in the infancy of genetic engineering and because of this will be the first mammalian species having all its genes characterized^13^. The fact that most of the laboratory strains of the house mouse used in research show aberrations in the circadian clock generators entrainment to earth’s solar day by photoperiodic rhythmicity of melatonin secretion and yet disclose circadian and infraradian biorhythmicities remains a challenge to investigate^16, 65^.

Although the driving engine of the slow oscillation in spontaneous activity observed here remains unknown, the large amplitude of the recurring variation in activity should have an impact at the system level. By using the feed consumption data available for 6 of the 14 cages studied, we found that feeding and activity level co-varied to a significant degree linking the metabolic state of the animals with the alternating base-line activity level (Fig. 4C). In this context it should be mentioned that observations on somatic variations in inflammatory responses, host-parasite interaction as well as epidemiology of ulcerous dermatitis common to the C57BL/6 strain, show distinct season-like variations during the year^77–79^ suggesting that circannual oscillations are not limited to behaviours in laboratory mice. Season-like variations in behaviours and somatic state of the commonly used C57Bl/6mouse have hitherto not received wide recognition but will need to be seriously considered in future experimental designs.

## Data files and data availability

All 14 data sets (one for each cage used) arranged in three groups (1-3) are available as three comma delimited csv files at: datadryad.org (doi:10.5061/dryad.n5tb2rbsf).

All data files have the following column headings for each cage:

Example from file group 1

| board | n_count | local_dt | age | AvgGlobal | group |
| --- | --- | --- | --- | --- | --- |
| A05 | 234 | 2016-11-29 00:00 | 49 | 0.013533 | grp1 |
| A05 | 238 | 2016-11-29 00:01 | 49 | 0.006303 | grp1 |

Where **board** is the cage ID, **n_count** is number of samples min^−1^, **local**_**dt** is local date followed by time stamp (CET), **age** is the age in days of the animals inside the cage, **AvgGlobal** is the average number of activations min^−1^ across all twelve electrodes, and **group** is the cage cohort that the cage belongs to.

The CSV files have been arranged to not violate the maximum number of input-lines (1 million lines) by certain applications, thus, data of each cage within each group have been listed in columns side by side.

Lights-on and Lights-off, respectively, happened every day at 04:00 (h:min; zeitgeber time 0) and 16:00 (h:min; zeitgeber time 12) dt (local).

## Acknowledgements

Help and advice from technical staff at the Wallenberg laboratory facility is gratefully acknowledged as is the use of equipment at the KI mouse clinic (KIMC). We are indebted to Art director Maria Larson for help with the art work of this study. We would like to thank Mara Rigamonti; Fabio Iannello and Giorgio Rosati for discussions on statistical calculations and help with access to the raw data.

## Authors contribution

KP: local management and execution of the experiments including collecting data, script writing, data analyses and preparation of illustrations, tables and statistical analyses. Editing of manuscript.

ER: Functional data analyses, preparation of illustrations and reading and commenting on the manuscript.

BU: study design and supervision. Drafting and editing of manuscript.

